# Feasibility of multimodal metabolic analysis for detecting early changes in acute neuroinflammation

**DOI:** 10.64898/2026.01.09.698663

**Authors:** James T. Grist, Ilia Evstafev, Dominika Olesova, Signe E. Nynäs, Matej Orešič, Alex M. Dickens, Damian J. Tyler, Yvonne Couch

## Abstract

Given the prevalence of metabolic perturbations in a variety of neurological and neurodegenerative diseases, understanding and monitoring brain metabolism is a key step in our advancement of therapies. The details of the citric acid cycle were established at the beginning of the last century but only recently have its metabolic intermediates been observed *in vivo* in the brain. In this study, we employed orthogonal analyses to investigate metabolic alterations in response to acute neuroinflammation *in vivo,* demonstrating a multi-technique approach that could be used for future studies.

Hyperpolarized ^13^C-pyruvate spectroscopy revealed an early decline in pyruvate metabolism via pyruvate dehydrogenase (PDH), leading to reduced ^13^C-bicarbonate formation. This metabolic disruption occurred despite the absence of structural or perfusion changes on conventional MRI. Further analysis of polar metabolites in the ipsilateral hemisphere confirmed ongoing inflammatory processes. These findings highlight the potential of this dual technique approach to inform upon metabolic changes due to neuroinflammation.

Combining methods to probe metabolism in invasive (metabolomics) and non-invasive (hyperpolarized MRI) manners, this represents a promising translational approach for real-time metabolic assessments in an area of the body, the brain, where studying processes such as metabolism has traditionally been challenging. This study has demonstrated the approach to monitor changes in metabolism in response to inflammation in the brain.

## Introduction

Current understanding suggests that ongoing inflammation in the brain is an important contributor to the development of neurodegenerative diseases such as dementia, but that it is also a significant factor in neuroinflammatory diseases such as multiple sclerosis (MS)(1–3). Monitoring on going inflammatory processes, and restoring homeostasis, is crucial for minimizing ongoing brain damage after injury, as well as monitoring the efficacy of anti-inflammatory therapies.

The readout of metabolic pathways in the brain may go some way to informing us of ongoing inflammatory pathology, as a close but indirect biomarker of the activated central nervous system (CNS) immune system (4). For example, astrocytes harness aerobic glycolysis to metabolise glucose to lactate, whereas quiescent microglia and neurons rely upon mitochondrial oxidative phosphorylation (OXPHOS) to fuel adenosine triphosphate (ATP) production through the Tricarboxylic Acid (TCA) Cycle (5). [1-^13^C]pyruvate can be metabolised through glycolysis (via lactate dehydrogenase (LDH), with [1-^13^C]lactate produced) and OXPHOS (via pyruvate dehydrogenase (PDH), hallmarked by ^13^C bicarbonate production). It is possible to monitor these pathways in the brain using hyperpolarized magnetic resonance imaging (MRI) (6, 7). Hyperpolarized carbon-13 MRI harnesses the transient increase in the available signal from a ^13^C-labelled substrate, often via the process of dynamic nuclear polarization, to allow for the real-time tracking of the metabolic fate of that label *in vivo* (7). Previous work has demonstrated that macrophage recruitment after cardiac ischemia can be monitored using [1-^13^C]pyruvate (8), as well as demonstrating increased [1-^13^C] lactate formation in models of neuroinflammation (9). One previous study has assessed the metabolic changes associated with lipopolysaccharide (LPS) in the mouse brain, detecting upregulated lactate production at 7 days post injection, correlated with markers of astroglyosis and microglial proliferation (10), demonstrating the potential for the detection of neuroinflammation behind the closed blood-brain barrier.

Clinically, the current technique for monitoring inflammation in the brain is positron-emission tomography. For example, mitochondrial translocator protein (TSPO) has been shown to be upregulated during periods of inflammation in a variety of cells, including astrocytes and microglia (11). By using radiolabelled ligands, such as ^11^C PK11195, it is possible to image this activity *in vivo* to study inflammation in brain injury (12). Areas of high TSPO binding have traditionally been thought to represent microglial activation in the brain. However, recent research has demonstrated that whilst this may be true in rodents, it remains unclear whether a similar pattern occurs in humans (13). Moreover, in some inflammatory diseases, such as MS, there is not a significant upregulation of TSPO and as such, use of this technique to monitor inflammation may be limited (14).

Metabolic imaging, as opposed to imaging specific inflammatory processes, can be carried out using ^18^F(lourine) Deoxy Glucose Positron Emission Tomography (^18^FDG-PET) or Magnetic Resonance Spectroscopy (MRS). FDG-PET uses radiolabelled deoxy glucose which cannot participate in the later stages of metabolism or exit the cell, resulting in images displaying increased signal intensity from metabolically active cells. However, neurodegenerative diseases such as dementia are known to result in metabolically underactive cells (15, 16) meaning that traditional methods such as PET are less sensitive to subtle changes in mitochondrial metabolism, and as such may fail to miss earlier stages of the neurodegenerative process. One of the earlier stages of neurodegeneration may be neuroinflammation (17).

A combined approach harnessing spatial metabolomics, hyperpolarized MR, molecular biology, and histopathology has been demonstrated in the oncological field (18), with results allowing a wide array of pathways to be explored and then a translational, tailored, ^13^C labelled biomarker derived for non-destructive experiments. These orthogonal techniques allow for holistic profiling of the tissue level response to an oncological challenge, and so the primary aim of this project was to translate this combined approach to a model of acute neuroinflammation. This proof-of-concept study will then pave the way for using this approach in more complex neuroinflammatory pathologies.

## Materials and Methods

### Animals

All procedures conformed to the Animal (Scientific Procedures) Act (1986) and were approved by the University of Oxford Animal Ethics Committee and the UK Home Office. The studies were conducted, and the manuscript prepared in accordance with the ARRIVE guidelines (19). All experimental procedures were carried out in the light phase of the animals’ light-dark cycle. Four adult male Wistar-Han rats were housed under standard light-dark conditions (12 hours) and allowed access to food and water *ad libitum*.

### Surgery

Stereotaxic procedures were carried out as previously described (20). Animals were anaesthetized using 2 % isoflurane in 60:40 oxygen:nitrous and mounted on a stereotaxic frame. The skull was exposed and a burr hole was drilled above the striatum (+1A/P, -2.5M/L, -4.5D/V from Bregma). 10 μg of lipopolysaccharide (*E.Coli* B26:06; Sigma-Aldrich, UK) in 1 μl of sterile 0.9 % saline was injected over a period of 5 minutes using a glass micro-needle. The needle remained in place for 2 minutes prior to slow withdrawal. Adsorbable sutures were used to close the wound. Animals were allowed to recover for 24 hours prior to imaging and tissue collection.

### Imaging

Animals were anesthetised with 2.5% isoflurane in 60:40 % oxygen: nitrous, a tail vein cannula inserted, and reduced to 2% when in the magnet as per a previously established protocol (21). Animals were placed in a custom-made cradle and placed inside of a 7 T magnet (Agilent magnet, Varian console) with a 2-channel ^13^C coil (Rapid Biomedical, Rimpar, Germany) used for reception and a ^1^H/^13^C volume coil (Rapid Biomedical) for transmit. Approximately 30 µL of [1-^13^C]pyruvate was prepared with 3 µL of 1:50 gadolinium in water (Dotarem, 1:50 dilution in H_2_O, 3uL:30uL diluted solution:[1-^13^C]pyruvate) and trityl radical (OXO63, 15 mM) and polarized as previously described (21). After approximately 1 hour of polarization, the sample was dissolved with 4.5 mL sodium hydroxide heated to approximately 175 degrees. Animals were injected with 1 mL of hyperpolarized [1-^13^C]pyruvate over 10 s, with 200 µL of saline used as a flush, and a 2-slice acquisition protocol used to acquire data from each hemisphere (Repetition time per slice = 500 ms, temporal resolution for both slices = 1 s, flip angle = 15 degrees, bandwidth = 5kHz, slice thickness = 10 mm). The ^13^C coil was replaced with a ^1^H 4-channel surface receive coil and T_1_-(pre and post gadolinium infusion, 3D Gradient and RF spoiled gradient echo, Field of View (FOV) = 40 mm^3^, matrix = 256x256x64, reconstruction matrix = 512x512x128, Repetition time (TR) = 5 ms, Echo Time (TE) = 1 ms, flip angle = 12 degrees) and perfusion weighted-imaging (200uL Dotarem infused over 0.5 seconds with 300uL saline flush, 2D Gradient Echo, FOV = 30 mm^2^, slice thickness = 1 mm, acquisition matrix = 128x64, reconstruction matrix = 128x128, TR = 20ms, TE = 10 ms, Temporal resolution = 1.28 s). Spectra were summed in the time domain and quantified using jMRIUI v5.2, with the fit for each metabolites used to calculate [1-^13^C]lactate: [1-^13^C]pyruvate, ^13^C bicarbonate: [1-^13^C]pyruvate, and ^13^C bicarbonate:[1-^13^C] lactate. Perfusion data were processed using model free deconvolution (21). T_1_ weighted imaging pre- and post-contrast injection were visually assessed for enhancement.

### Tissue Processing

Animals were intracardially perfused with cold 0.9% saline containing 10 U/ml of heparin. Brains were snap frozen using a dry ice/isopentane bath and stored at - 80°C until processing. 12 μm sections were cut on a cryostat and serial sections were taken onto SuperfrostPlus slides for immunohistochemistry, as well as being split into ipsilateral and contralateral hemispheres for RNA and protein extraction. Slides and frozen tissue were also stored at -80°C until further use.

### qPCR

RNA was extracted from approximately 25 mg of frozen brain tissue using the Qiagen© RNEasy Mini Kit with QiaShredders. RNA concentration was measured using a NanoDrop (ThermoFisher) and 800 ng of sample was converted to cDNA using the Applied Biosystems High Capacity cDNA conversion kit. Real-time qPCR was performed with duplicates for each sample (15 ng/well) using SYBR green qPCR master mix (PrimerDesign) and the Applied Biosystems QuantStudio Flex 7 with the following cycle conditions: hot start 2 m at 95°C, 40 cycles of 15 s at 95°C and 60 s at 60°C with data acquisition during the 60°C phase. The following primers were purchased from Merck as Pure&Simple Primers: *IL-1*β: F: CACCTCTCAAGCAGAGCACAG; R: GGTTCCATGGTGAAGTCAAC; *IL-6*: F: TCCTACCCCAACTTCCAATGCTC; R: TTGGATGGTCTTGGTCCTTAGCC. *ICAM*: F: AAACGGGAGATGAATGGTACCTAC; R: TGCACGTCCCTGGTGATACTC; *VCAM*: F: GGCTCGTACACCATCCGC; R: CGGTTTTCGATTCACACTCGT; *PDK1*: F: TACAGAACCAACCACGAGGC R: CCACATTTGGCTTTGCCAGG *PDK2*: F: ACCCAGTCTCCAACCAGAAC; R: GAGATGCGGCTGAGGTAGAA. *PDK3*: F: TCCTAGCGCTCTTGTACCCT; R: CACCCAAGTACCACACCTCC*PDK4*: F: GGATTACTGACCGCCTCTTTAGTT; R: GCATTCCGTGAATTGTCCATC GAPDH: F: GCAAGTTCAACGGCACAG r: CGCCAGTAGACTCCACGAC Relative expression was determined using the Pfaffle method (22) and normalized to the housekeeping gene GAPDH and further normalized to the contralateral hemisphere of the control animals.

### Western Blotting

Brain tissue was homogenized in ice-cold lysis buffer (50 mM Tris-HCl, pH 7.4, 150 mM NaCl, 1% Triton X-100, 1% sodium deoxycholate, 0.1% SDS) supplemented with a protease and phosphatase inhibitor cocktail (Roche, Basel, Switzerland). The homogenates were centrifuged at 12,000 × g for 15 minutes at 4°C, and the supernatants were collected for protein quantification. Protein concentrations were determined using the bicinchoninic acid (BCA) assay (Thermo Fisher Scientific, Waltham, MA, USA). For each sample, 10 µg of protein was mixed with 4× NuPAGE LDS sample buffer (Invitrogen, Carlsbad, CA, USA) and 10× reducing agent (β-mercaptoethanol; Invitrogen) and heated at 70°C for 10 minutes. Proteins were separated on 4–12% Bis-Tris gradient gels (NuPAGE, Invitrogen) using MOPS SDS running buffer (NuPAGE, Invitrogen) under constant voltage (200 V) for approximately 50 minutes. Following electrophoresis, proteins were transferred onto polyvinylidene difluoride (PVDF) membranes (0.45 µm pore size; Millipore, Burlington, MA, USA) using a semi-wet system according to the manufacturer’s protocol (Bolt^TM^ Transfer Buffer). The membranes were blocked in 5% non-fat dry milk (w/v) prepared in Tris-buffered saline with 0.1% Tween-20 (TBST) for 1 hour at room temperature. The membranes were incubated overnight at 4°C with primary antibodies diluted in 5% non-fat dry milk (w/v) in 0.1% TBST. Primary antibodies were against HSP60 (housekeeping), pyruvate dehydrogenase (PDH), phosphorylated PDH (phPDH) and pyruvate kinase M (PKM). All antibodies were purchased from AbCam (Cambridge, UK) and used at 1:5000. After three washes with TBST, membranes were incubated with HRP-conjugated secondary antibodies (1:20,000) for 1 hour at room temperature. Signal detection was performed using an enhanced chemiluminescence (ECL) reagent (Bio-Rad, Hercules, CA, USA) and imaged with a ChemiDoc Imaging System (Bio-Rad). Densitometric analysis was performed using Bio-Rad ImageLab software, and target protein signals were normalized to the corresponding loading control (HSP60) or as a ratio of phPDH:PDH.

### PDH Activity Assay

PDH activity was determined using a commercially available PDH activity assay kit which was run according to the manufacturers instructions (AbCam, Cambridge, UK). Briefly, tissue was weighed and homogenized in 10x volume tris buffered saline made with 10mM NaF and 1mM PMSF (phenylmethylsulfonyl fluoride used to inhibit serine proteases). Protein was quantified using BCA assay. Protein was normalised to 2.5mg/ml using TBS + PMSF and detergent added at 1:20. Samples were incubated with some shaking on ice for 15 minutes before centrifugation at 1000g for 10 minutes. 150 µl per well of the precoated assay plate was added to 50 µl of PDH assay buffer and incubated overnight at 4°C. Detection buffer was added immediately prior to running on a colourimetric plate reader running at 37°C for 45 cycles.

### Metabolomics Sample Preparation

Rat plasma was used as sample matrix. For polar metabolites and lipid analysis, 40 µL of plasma was mixed with 480 µL of isotopically labelled standard mixture in MeOH:MTBE:IPA (20:15:15, v/v). Samples were vortexed for 10 minutes, followed by incubation at -20°C for 45 minutes. Protein removal was conducted using a protein precipitation plate (55263-U, Supelco), and 100 µL of the filtrate was transferred to glass insert vials. The aliquots were evaporated and reconstituted in 50 µL water for polar metabolites analysis. The leftover filtrate was used for the lipidomics assay.

Working solutions were prepared at 9 concentration levels by serial dilution from 0.125 µg/mL to 25 µg/mL in water and from 0.039 µg/mL to 10 µg/mL in chlorophorm:methanol (2:1, v/v) for polar metabolites and lipids respectively. The lipidomics master mix contained CE(18:1(9Z)), CE(18:2(9Z, 12Z)), Cer(d18:0/18:1(9Z)), Cer(d18:1/18:1(9Z)), LysoPC(18:0), LysoPC(18:1), LysoPE(18:1), PC(16:0/16:0), PC(16:0/18:1), PC(18:0/18:0), PE(16:0/18:1), TG(18:0/18:0/18:0), C16 dihydroceramide (d18:0/16:0), TG(16:0/16:0/16:0). The polar metabolites master mix contained glucose 1-phosphate, glucose 6 phosphate, fructose 6 phosphate, dihydroxyacetone phosphate, fructose 1,6 biphosphate, glyceraldehyde 3 phosphate, 2,3-diphospho-D-glyceric acid, D-2-phosphoglyceric acid, PEP/phosphpenopyruvate, lactate, glycerol-3-phosphate, pyruvate, 6-phosphogluconic acid, D-ribose 5-phosphate, acetyl coA, citrate, alpha-ketoglutarate, succinyl-CoA, succinate, fumarate, malonyl coenzyme A, malate, Itaconate, glutarate, isocitrate, oxaloacetic acid, L-arginine, glycine, alanine, valine, leucine, L-lysine, L-trytophan, L-phenylalanine, GABA, isoleucine, histidine, 3-hydroybutyric acid, L-proline, cysteinie, octanoic acid, 2-oxo-glutarate, methionine, L-threonine, L-serine, aspargine, phenylpyruvic acid, arginine, ketoleucine, 2-OH-butyric acid, taurine, omithine, ascorbic acid, creatine, xylulose-5-phosphate, erythrose-4-phosphate, sedoheptulose 7-phosphate, adenosine 5′-monophosphate.

Calibration curve samples were prepared by combining 10 µL of corresponding polar metabolites working solution*s* and 10 µL of lipid working solution, along with 240 µL of isotopically labeled internal standard mixture comprising of succinic acid-d4, valine-d8, glutamic acid-d5, glutamine-13C5, glucose-13C6, GABA-d6, glycerol-d5, indole-3-acetic acid-d5, tyrosine-d4, stearic acid d35, melatonin-d4, nicotinamide-d4, hippuric acid-d5, 18:1-d7 LYSO PE, 15:0-18:1-d7 DG, 16:0 cholesteryl-d7 ester, 18:1-d9 SM, 15:0-18:1-d7-15:0 TG, 15:0-18:1-d7-PE, 15:0-18:1-d7-PC, 18:1-d7 LYSO PC, C15 ceramide-d7, C13-dihydroceramide-d7 at 1 µg/mL concentration each in MeOH:MTBE:IPA (20:15:15, v/v).

To track analytical performance a set of solvent QC samples and pooled matrix QC samples were prepared and were injected after every 10 samples injections of the study samples. The solvent QC samples prepared out of level 6 working solution *(3.125* µg/mL for polar metabolite in the mix and 1.25 µg/mL for the lipids) according to the calibration curve samples procedure. The pooled matrix QC samples were prepared by pooling 20 µL of all samples, vortexing, and aliquoting into 40 µL portions for subsequent preparation.

### Liquid Chromatography

#### Polar metabolites analysis

Liquid chromatography was performed on an ExionLC AD system, comprising two pumps, an autosampler, an AC column oven, and a system controller. A Waters Atlantis Premier BEH C18 AX column (1.7 µm, 2.1 × 100 mm) was used for separation. Mobile phases were water with 0.1% formic acid (v/v) and 10 mM ammonium formate (Phase A) and methanol with 0.1% formic acid (v/v) and 10 mM ammonium formate (Phase B). The gradient profile was as follows: 0–2 min, 100% A; 2–4 min, linear ramp to 100% B; 4–10 min, 100% B; 10.1–16 min, 100% A. The flow rate was maintained at 400 µL/min, with the column and autosampler temperatures set to 50°C and 15°C, respectively. Injection volume was 10 µL, and the total run time was 16 minutes.

#### Lipid analysis

Liquid chromatography was performed on an ACQUITY Premier UPLC system, comprising binary solvent manager, a <u>sample manager and</u> a column <u>manager</u>. A Waters ACQUITY BEH C18 column (1.7 µm, 2.1 mm × 100 mm) was used for separation. Mobile phases were 10mM of ammonium acetate in water with 0.1% formic acid (v/v) (Phase A) and 10mM of ammonium acetate in mixture acetonitrile:isopropanol (1/1, v/v) with 0.1% formic acid (v/v) (Phase B). The gradient profile was as follows: 0–2 min linear ramp 35% B to 80% B; 2–7 min, linear ramp to 100% B; 7–14 min, 100% B; 14-14.1 linear ramp to 35%B; 14.1–<u>21</u> min, <u>35</u>% B. The flow rate was maintained at 400 µL/min, with the column and autosampler temperatures set to 50°C and 15°C, respectively. Injection volume was 1 µL, and the total run time was <u>21</u> minutes.

### Mass Spectrometry

#### Polar metabolites analysis

A SCIEX QTOF 6600 equipped with a Duospray ion source was employed for mass spectrometry. Ionization was performed in negative mode with the following source parameters: source temperature, 650°C; spray voltage, -4500 V; ion source gas 1 and gas 2, 40 psi each; curtain gas, 25 psi; declustering potential (DP), -40 V; collision energy (CE), -10 V; and entrance potential (EP), -10 V. MS acquisition utilized an accumulation time of 0.4 seconds per scan over a mass range of 50–1000 m/z. The MS/MS data was obtained by information dependent acquisition (IDA) and was performed with a survey scan accumulation time of 0.25 seconds, accumulation time for MS/MS automatically selected precursor scan was set to 0.1 seconds, and a precursor mass tolerance of 50 ppm. Up to four MS/MS spectra were collected per cycle.

#### Lipid analysis

A SCIEX QTOF 7600 equipped with a OptiFlow (50-200µL probe) ion source was employed for mass spectrometry. Ionization was performed in positive mode with the following source parameters: source temperature, 300°C; spray voltage, 5500 V; ion source gas 1 and gas 2, 70 psi each; curtain gas, 50 psi; declustering potential (DP), 80 V; collision energy (CE), 10 V. MS acquisition utilized an accumulation time of 0.25 seconds per scan over a mass range of 100–1000 m/z. The MS/MS data was obtained by information dependent acquisition (IDA) and was performed with a survey scan accumulation time of 0.1 seconds, accumulation time for MS/MS automatically selected precursor scan was set to 0.1 seconds, and a precursor mass tolerance of 50 mDa. Up to ten MS/MS spectra were collected per cycle.

### Mass Spectrometry Imaging sample preparation

The MALDI matrix was applied by sublimation. Super-DHB (Sigma-Aldrich) was used as a matrix at a 40 mg/mL concentration in acetone. The vacuum for the sublimation was created by MD 1C +AK+EK chemistry vacuum system (VACUUBRAND GMBH). A volume of 750 µL was applied on the bottom of the sublimator chamber, dried with nitrogen and submitted to the vacuum with installed sample slide held above on an ice finger, the sublimation was carried out for 15 minutes under 140°C.

### Mass Spectrometry Imaging Analysis

#### Small Molecules

Samples were measured on Bruker timsTOF fleX equipped with a post-ionisation laser (Bruker Daltonics) in positive mode from m/z 50 to 650 with 28 lasershots per pixel using a 20 µm laser settings (20 µm spot size; 20 µm step size). The ion mobility separation was set for 100 ms, 1/K0 was ranged from 0.4 to 1.1 V·s/cm^2^.

#### Lipids

Samples were measured on Bruker timsTOF fleX equipped with a postionisation laser (Bruker Daltonics) in positive mode from m/z 300 to 1350 with 40 lasershots per pixel using a 20 µm laser settings (20 µm spot size; 20 µm step size). The ion mobility separation was set for 200 ms, 1/K0 was ranged from 0.7 to 1.8 V·s/cm^2^.

### Mass Spectrometry Data Treatment

All imaging experiments were loaded into SCiLS Lab (v2023bPro, Bruker Daltonics) using the default settings with the parameters from FlexImaging. The data were root mean square (RMS) normalized, and a list of features was generated and aligned in SCiLs Lab. The compounds that show a difference in distribution between hemispheres were found using a Receiver Operating Characteristic (ROC) value by comparing manually drawn striatum regions. ROC values were calculated for each tissue section. The intensities of the ion images are individually adjusted for the sections to best show the differences between hemispheres.

All the features underwent filtering based on acquired ROC values. Features with higher than 20% CV for case and 15% for control samples were filtered out. The obtained list was compared case mean ROC vs control mean ROC. The features with a difference of less than 0.05 were filtered out. Next, features were filtered based on case accuracy, calculated as the mean of all control ROC AUC values, normalized to a 0.5 baseline and multiplied by 100. Features with an accuracy between 90% and 110% were retained. 4 features were selected for further structure confirmation by MS/MS spectra, which were acquired after the data treatment by utilising the incurred samples.

### Statistics

For imaging studies, metabolic ratios were compared between hemispheres using the Wilcoxon Rank Sum test. For all other analysis standard statistical tests (ANOVA, t-test), with adjustment for normality, were performed in Prism 10. A p-value of <0.05 was considered significant.

Lipidomics and metabolomics data were processed and statistically evaluated in R (https://www.r-project.org, 2025, v4.5.0) using an in-house developed script. The data were transformed by natural logarithm (ln), and the mean centering was applied. Data were evaluated using multivariate statistical analysis (principal component analysis, PCA). The univariate statistical analysis was based on the parametric t-test combined with the log2 fold-change (log2 ratio of means) and is provided in supplementary Tables 1 & 2. Statistically significant features (p-value < 0.05) are presented in form of a heatmap. Spearman’s R was calculated using Hmisc package and visualized as a heatmap. Functional analysis of untargeted metabolomics data was performed using Metaboanalyst v6.0 (https://www.metaboanalyst.ca) with mass tolerance 10 ppm.

## Results

### Intrastriatal LPS injection results in ipsilateral neuroinflammation

Our first aim was to establish the efficacy of an acute model of neuronflammation in terms of histology and molecular biology. Histologically, animals showed an increase in markers of immune cell and vascular activation with all data showing a significant main effect of LPS and an interaction between inflammation (LPS) and hemisphere. Increased ICAM staining in the ipsilateral hemisphere of LPS injected animals (two-way ANOVA; Sidak’s p<0.01; Fig.1A & E).

**Figure 1.**
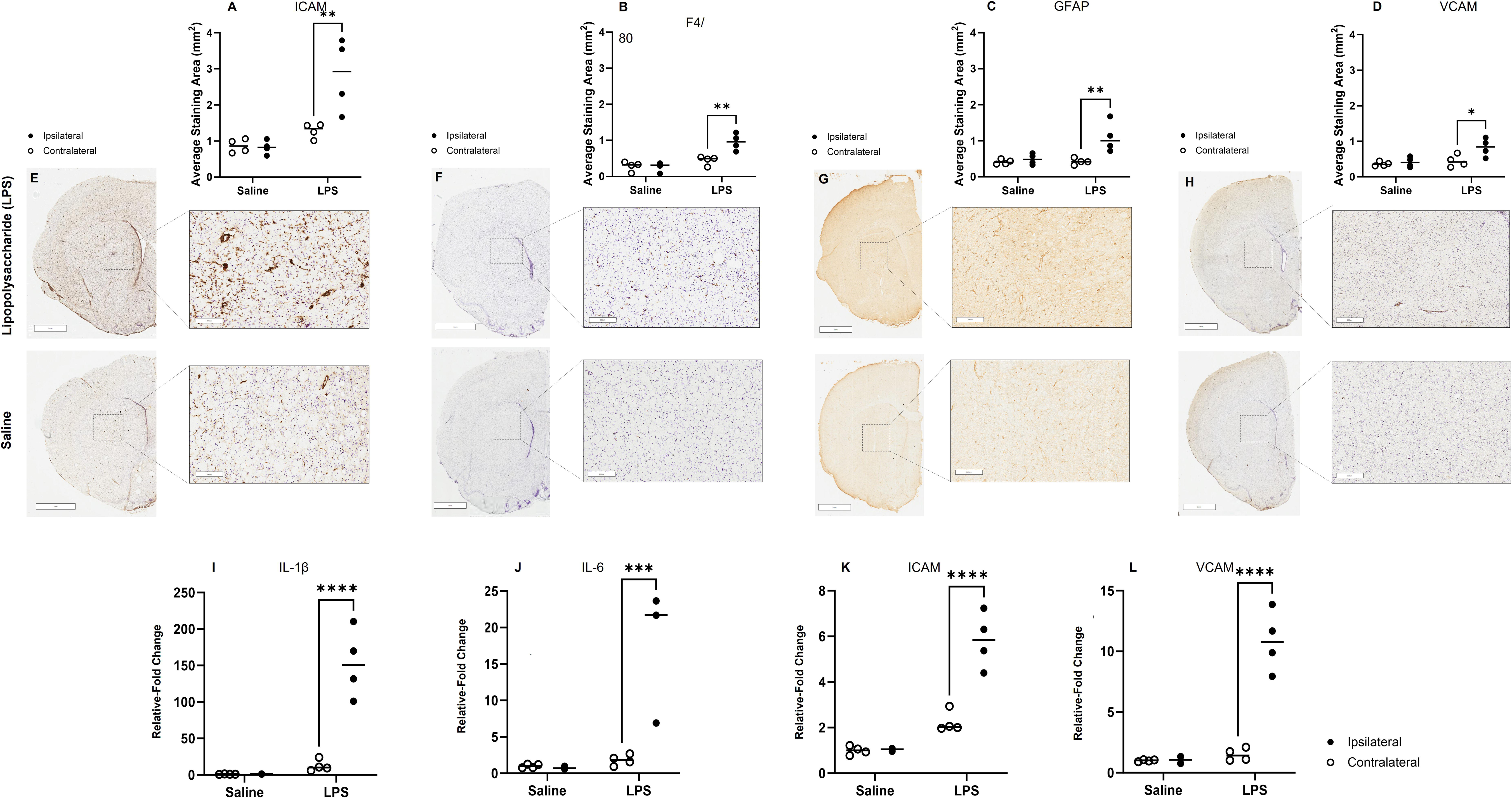
LPS-induced neuroinflammation increases markers of immune and vascular activation in the ipsilateral hemisphere. Representative images and quantification show increased immunoreactivity for ICAM (A, E), F4/80 (B, F), GFAP (C, G), and VCAM (D, H) in the ipsilateral hemisphere following LPS injection, indicating activation of endothelial cells, microglia/macrophages, astrocytes, and vascular endothelium respectively. Quantitative PCR demonstrated significant increases in mRNA expression of IL-1 (I), IL-6 (J), ICAM (K), and VCAM (L), with a main effect of inflammation, hemisphere, and a significant interaction between the two factors (two-way ANOVA; IL-1 and ICAM: inflammation – p<0.01, hemisphere – p<0.001, interaction – p<0.01; IL-6: inflammation – p<0.01, hemisphere – p<0.01, interaction – p<0.01; VCAM: p<0.001 for all effects). Post-hoc testing revealed a significant increase in gene expression in the ipsilateral hemisphere of LPS-injected animals for IL-1 (Sidak’s; p<0.0001; A), ICAM (Sidak’s; p<0.0001; C), IL-6 (Sidak’s; p<0.0001), and VCAM (Sidak’s; p<0.001). Data are presented as mean ± SEM; n = 4 per group. Scale bar on large image represents 2mm and on the inset panels, 200µm.

F4/80 expression levels were significantly increased in the ipsilateral hemisphere after LPS injection, reflecting increased microglia/macrophage activity (two-way ANOVA; Sidak’s p<0.01; Fig.1B & F). GFAP expression increased, reflecting activation of local astrocyte populations (two-way ANOVA, Sidak’s p<0.01; Fig.1C & G). Finally, VCAM expression levels were modest compared to ICAM but were significantly increased in the ipsilateral hemisphere of LPS-injected animals (two-way ANOVA; Sidak’s p<0.05; Fig.1D & H). For most inflammatory genes, our previous research has demonstrated that mRNA and protein expression are tightly correlated(23, 24) and as such, mRNA expression profiles are a good proxy for the inflammatory landscape in the brain.

IL-1 (Fig.1I) and ICAM (Fig.1K) showed a main effect of inflammation, a main effect of the hemisphere studied and an interaction between the effects (two-way ANOVA; inflammation – p<0.01, hemisphere – p<0.001, interaction – p<0.01 for both Fig.1I and Fig.1K). Post-hoc analysis revealed a significant increase in IL-1 (Sidak’s; p<0.0001; Fig.1I) and ICAM expression (p<0.0001; Fig.1K) in the ipsilateral hemisphere after LPS injection. In terms of IL-6 mRNA expression, there was a significant main effect of inflammation (two-way ANOVA; p<0.01; Fig.1J), a significant main effect of brain hemisphere (p<0.01) and a significant interaction between the effects (p<0.01). Post-hoc testing revealed an increase in IL-6 expression in the ipsilateral hemisphere in response to LPS (Sidak’s; p<0.0001). Finally, VCAM expression levels changed in response to inflammation (two-way ANOVA; p<0.001; Fig.1L), in different brain hemispheres (p<0.001) and there was an interaction between the effects (p<0.001). As with the other inflammatory genes, post-hoc testing showed an increase in the ipsilateral hemisphere in response to an injection of LPS (Sidak’s; p<0.001; Fig.1L).

### Acute neuroinflammation results in decreased bicarbonate:pyruvate and bicarbonate:lactate in the absence of structural changes

As well as metabolites used for biosynthesis, it was also important to use methods for studying metabolites used for energy production. Using perfusion weighted imaging and gadolinium we determined there was no significant difference in cerebral blood flow after an intrastriatal injection of LPS (t-test: p=0.6; Fig. 2A & B) nor was there any significant blood brain barrier breakdown (Fig.2C & D; pre- and post-gadolinium, respectively). The signal from injection of [1-^13^C]pyruvate was analysed in the ipsilateral and contralateral hemispheres (Fig.2E). There was no significant change in the [1-^13^C]lactate:[1-^13^C]pyruvate ratio between hemispheres, no difference between LPS and control groups and no interaction between main effects (two-way ANOVA inflammation: p=0.86; brain hemisphere p=0.99; interaction p=0.99; Fig.2F). There was a decrease in ^13^C bicarbonate:[1-^13^C]pyruvate ratio which was statistically significant in terms of inflammation (two-way ANOVA; p<0.05; Fig.2G) and brain hemisphere (p<0.01) and there was a significant interaction between inflammation and brain hemisphere (p<0.01). Post-hoc testing revealed a significant difference in the ratio of bicarbonate to pyruvate between contralateral and ipsilateral hemispheres in the LPS injected animals (Sidak’s; p<0.001).

**Figure 2.**
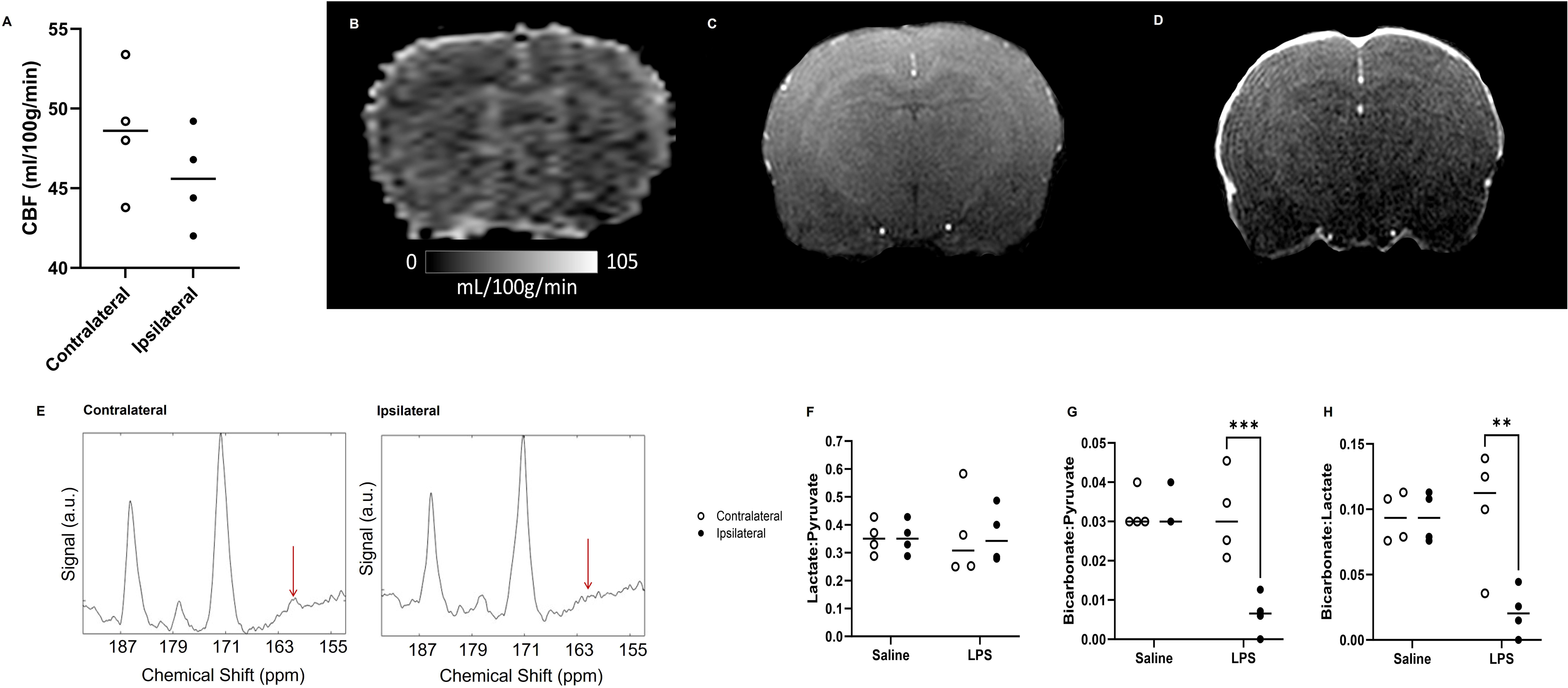
Assessment of cerebral perfusion and pyruvate metabolism following LPS injection. There was no significant difference in cerebral blood flow between groups (A; t-test; p = 0.6), with an example CBF map from an LPS brain shown in B. Blood-brain barrier integrity as assessed using pre- and post-injection of gadolinium contrast (C and D, respectively) showed no discernible contrast agent extravasation. Example hyperpolarized spectra from the ipsilateral hemisphere in the LPS brain showed a decrease in ^13^C bicarbonate formation (E) compared to contralateral (F). No significant difference in [1-13C] lactate: [1-13C] pyruvate was observed (F, p > 0.86), however a significant main effect of inflammation (p < 0.05), hemisphere (p < 0.01), and interaction (p < 0.01) was observed for the ^13^C bicarbonate: [1-13C] pyruvate ratio (G). For the ^13^C bicarbonate: [1-13C] lactate ratio, there was a main effect of hemisphere (p < 0.05) and interaction (p < 0.01), but not of inflammation (p = 0.06) (H). Post-hoc testing revealed significant differences between contralateral and ipsilateral hemispheres in LPS-injected animals (Sidak’s; G: p < 0.001; H: p < 0.01). Data are presented as mean ± SEM; n = 4 per group.

Similarly, ^13^C bicarbonate:[1-^13^C]lactate ratio was significantly different in different brain hemispheres (two-way ANOVA; p<0.05; Fig.2H) and there was an interaction between inflammation and brain region (p<0.01) but inflammation per se was not a significant main effect (p=0.06). Post-hoc testing revealed a significant difference in the ratio of bicarbonate to lactate between contralateral and ipsilateral hemispheres in the LPS injected animals (Sidak’s; p<0.01).

### Acute neuroinflammation changes the peripheral metabolite profile

We next measured the lipids and metabolites in the plasma of the LPS injected rats. These showed clear separation on a PCA plot (Fig.3A). There were also significant differences in metabolite and lipid levels (Fig.3B). Given the changes in bicarbonate observed in the brain via hyperpolarised imaging we next sought to identify if this was reflected in the peripheral metabolome. Therefore, we quantified pyruvate and lactate in the blood and then saw what metabolites and lipids correlated to the blood levels of lactate and pyruvate (Fig. 3C, Supplementary Tables 1 & 2). Functional analysis of metabolomics data showed glutathione and pyrimidine metabolism as most implicated in peripheral response to central neuroinflammation (Fig 3D).

**Figure 3.**
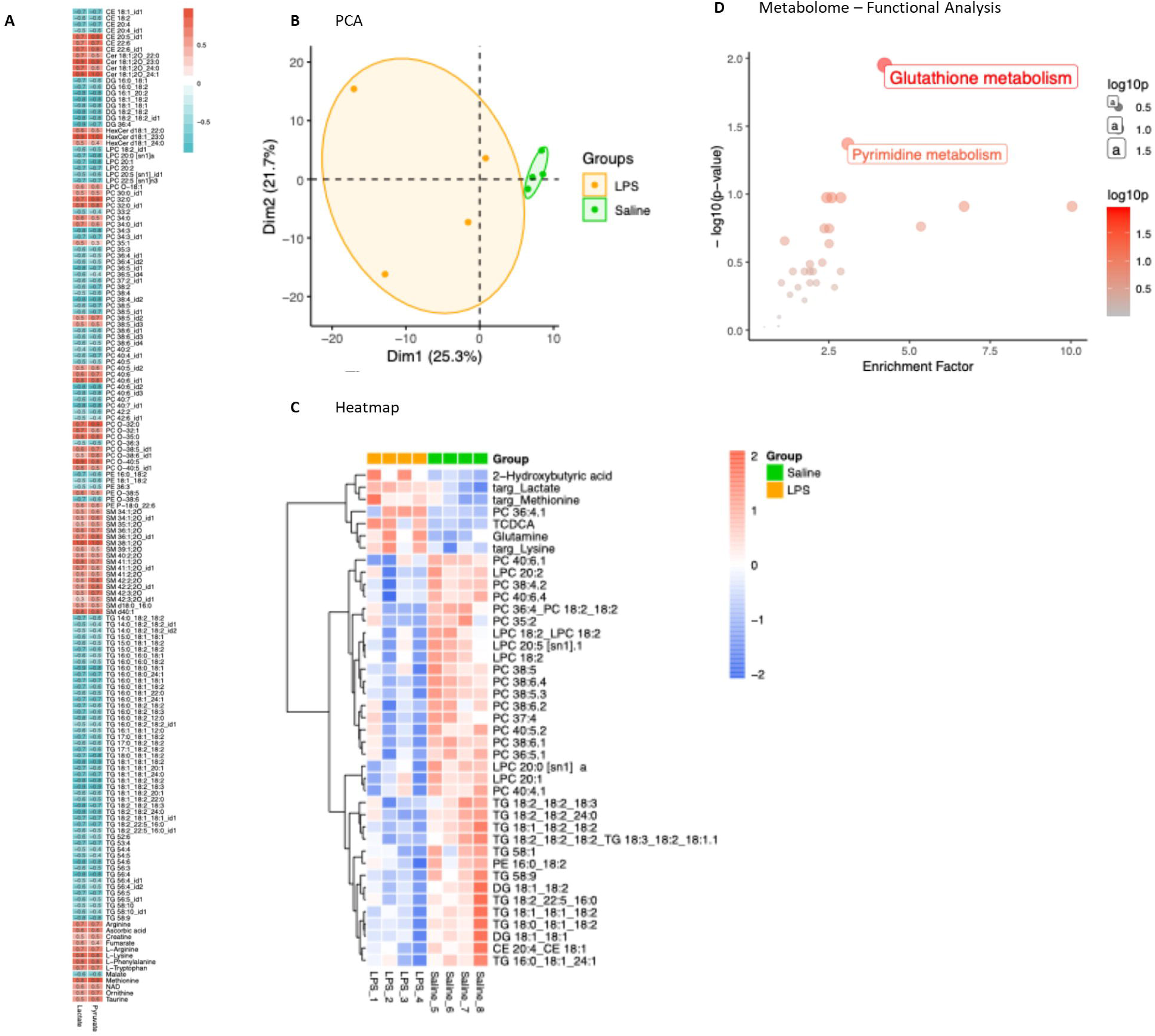
Plasma metabolome alterations are dominated by elevated glycerophospholipids and triacylglycerols. Spearman correlation heatmap (|R| > 0.5) showing metabolites associated with plasma lactate and pyruvate levels (A). Principal Component Analysis scores plot of plasma metabolome and lipidome (PC1 and PC2 explain 25.3% and 21.7% of variance, respectively) (B) Heatmap of significantly altered metabolites and lipids between LPS and Saline groups (Student’s t-test, p-value < 0.05) (C) Functional pathway analysis performed in MetaboAnalyst using the full plasma metabolome (m/z features) with associated p-values. (D).

### Acute neuroinflammation changes metabolism in the ipsilateral hemisphere

Spatial metabolomics was employed to investigate inflammation in the brain, as it enables the localization and characterization of metabolic alterations within specific anatomical regions, providing insight into region-specific biochemical responses to neuroinflammatory processes. Here, we demonstrated that metabolic reprogramming during inflammation results in ipsilateral changes in a variety of metabolites.

The obtained spatial metabolomics data showed several left right differences. The top four features as identified via the left right difference were: Cyclidine 2’3’-cyclic phosphate (Fig.4A-C), Glutathione (Fig.4D-F), N-palmitoyl phenylalanine (Fig.4G-I)and Prostaglandin (Fig.4J-L). The figure shows representative brain sections of rats injected with LPS (Fig. 4A, 4D, 4G, 4J) and brain sections of control rats injected with vehicle (Fig. 4B, 4E, 4H, 4K). In addition, data on the signal intensity in the striatum are shown in the corresponding brain sections (Fig. 4C, 4F, 4I, 4L). For the images we can clearly see an upregulation of all 4 molecules in the ipsilateral striatum compared to the contralateral and control animals. ROC values were calculated for the aforementioned compounds in rat brains. For each component, ROC values were determined in both the control and case samples. For cyclidine 2’3’-cyclic phosphate, the ROC was 0.515 in the control sample and 0.624 in the case sample. For glutathione, the ROC was 0.531 in the control sample and 0.634 in the case sample. For N-palmitoyl phenylalanine, the ROC was 0.481 in the control sample and 0.650 in the case sample. For prostaglandin, the ROC was 0.459 in the control sample and 0.637 in the case sample. The ROC values for all the samples are shown in the supplementary table (Supplementary Table 3).

**Figure 4.**
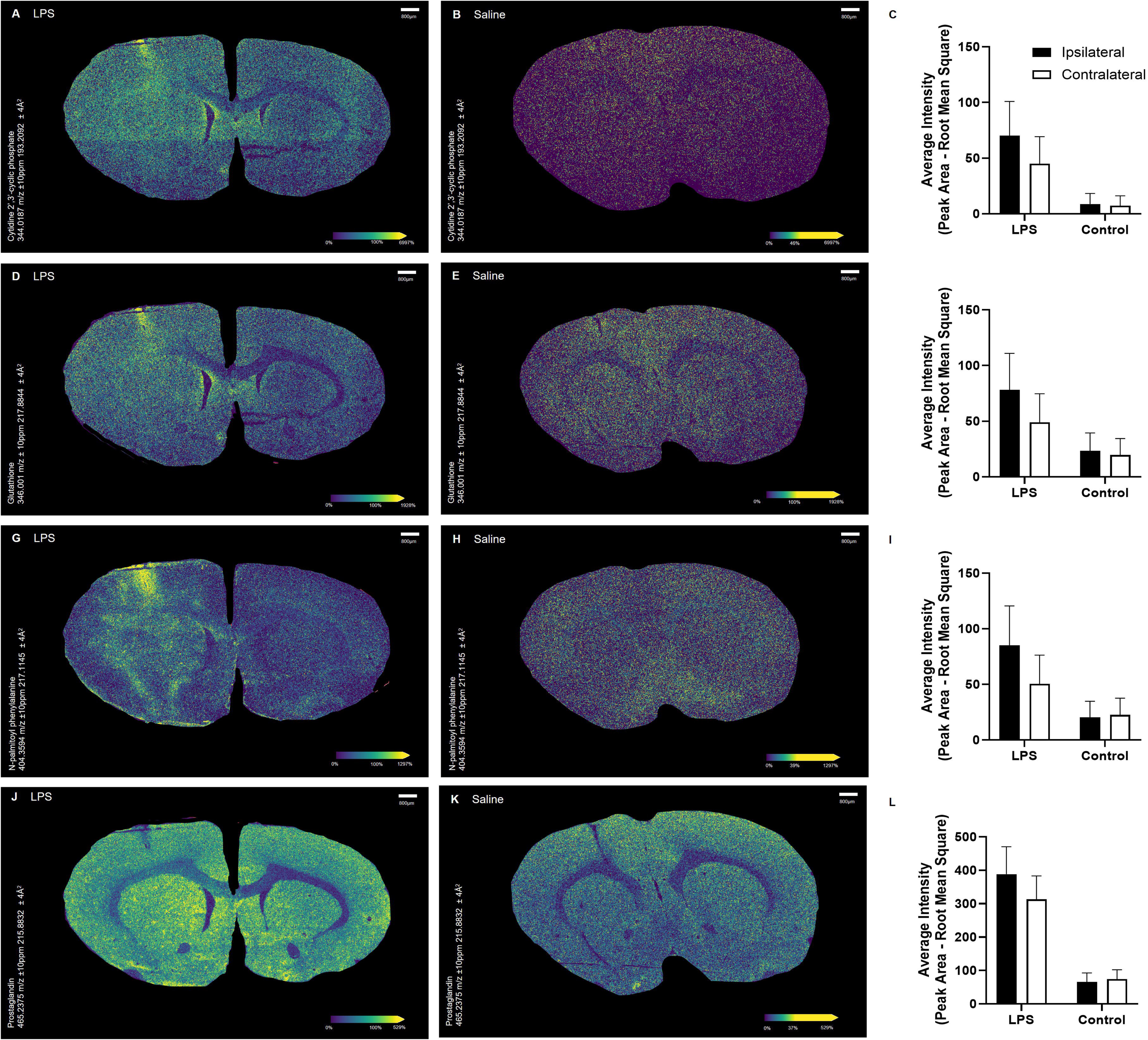
LPS-induced neuroinflammation results in ipsilateral changes in selected metabolites. Representative images are high spatial resolution (20µm) root mean square normalized ion images of LPS and saline injected brains for cytidine-2’3-cyclic phosphate (A & B), glutathione (D & E), N-palmitoyl phenylalanine (G & H) and prostaglandin (J & K). The brightness scale is adjusted sample-wise to assess sample to sample intensity variations. Graphs show average distribution of intensities of selected peak after root mean square normalization within the striatum (defined ROI) for cytidine-2’3-cyclic phosphate (C), glutathione (F), N-palmitoyl phenylalanine (I) and prostaglandin (L). Data are presented as mean ± SEM. Scale bar on representative images shows 800µm.

### Changes in energy metabolism measurements are accompanied by a trend in changes in pyruvate dehydrogenase activity

To discover whether changes in the metabolism of pyruvate might be associated with changes in pyruvate dehydrogenase, mRNA measurements of PDK isoforms, Western blotting and activity assays were performed.

PDK is known to regulate the activity of PDH and, as such, expression levels of mRNA are likely to change when the bioenergetic environment changes. Here, we found a significant main effect of inflammation on *PDK1* and *PDK2* mRNA expression (two-way ANOVA; p<0.0001 for both; Fig.5A&B) but no main effect of brain hemisphere (p=0.2, p=0.3; respectively) or any interaction between the main effects (p=0.09 for both). Post-hoc testing revealed a significant increase in both *PDK1* and *PDK2* expression in the ipsilateral hemisphere after LPS injection (Sidak’s; p<0.0001; Fig.5A&B). We found no significant effect of either brain hemisphere or inflammation, nor any interaction between the main effects in the expression levels of *PDK3* or *PDK4* (Fig.5C&D). Blotting (Fig.5E) showed a change in phPDH:PDH ratio in the ipsilateral hemisphere of LPS injected animals which did not reach significance, likely due to low statistical power (Fig.5F, p=0.06). To determine whether there was generic dysregulation of energy producing enzymes we also measured changes in PKM in the same tissue and found no significant differences in PKM expression between LPS and sham animals (Fig.5G). Finally, to determine whether the changes in ratio of phPDH to PDH reflected actual changes in activity, we used an activity assay to show a relative decrease in PDH activity in the ipsilateral hemisphere of LPS injected animals (Fig.5H). However, much like the PDH ratio there was a lack of power to this experiment (Fig.5H, p=0.06).

**Figure 5.**
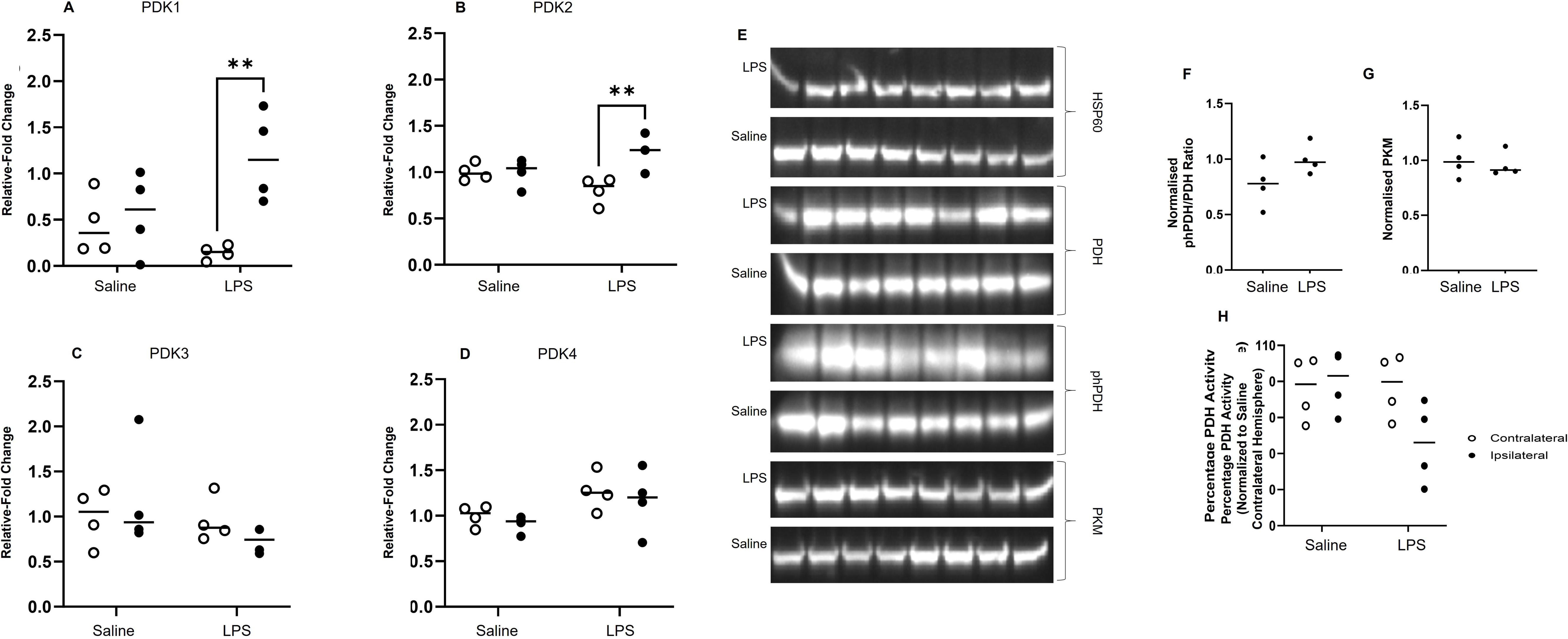
Inflammation-associated changes in pyruvate metabolism are accompanied by altered PDK1 and 2 mRNA expression and trends in PDH regulation. mRNA expression of PDK isoforms was measured in ipsilateral and contralateral brain hemispheres. Significant changes were observed in *PDK1* (A) and *PDK2* (B) but not *PDK3* (C) or *PDK4* (D). Both *PDK1* and *PDK2* mRNA expression showed a significant main effect of inflammation (two-way ANOVA; *p* < 0.0001) with increased expression in the ipsilateral hemisphere of LPS-injected animals (Sidak’s; *p* < 0.01; A & B). Western blotting revealed a non-significant trend toward increased phPDH:PDH ratio in LPS-injected animals (E, F; *p* = 0.06). PKM expression was unchanged between groups (G). PDH activity assay showed a trend toward reduced enzymatic activity in the ipsilateral hemisphere following LPS injection (H; *p* = 0.06). Data are presented as mean ± SEM; *n* = 4 per group.

## Discussion

This study used a combination of advanced technologies to probe the metabolic environment of the brain after acute LPS challenge. As a feasibility and proof-of-concept study with a necessarily small cohort, our aim was not to deliver definitive mechanistic conclusions but rather to demonstrate that these multimodal techniques can sensitively detect early metabolic consequences of acute neuroinflammation. The model was characterised through histology and enzymatic assays, revealing a prototypical pro-inflammatory response, demonstrated through increases in vascular reactivity and microglial and astrocyte activity, as well as upregulation of cytokine and chemokine mRNA.

Whilst the central injection of LPS has often been considered a reductionist way of studying neuroinflammation, it is being increasingly used as a way of investigating how neuroinflammation might precipitate diseases such as Alzheimer’s and Parkinson’s (25). However, in the context of a pilot-style study such as ours, this reductionism is an advantage, allowing us to isolate early metabolic effects with fewer confounding variables. In fact, we know that inflammation often precedes physical and cognitive impairment in a variety of neurodegenerative diseases (26, 27). Novel therapies targeting inflammation (28) and metabolic processes (29) are becoming increasingly in vogue and, as such, it will become ever more important to be able to monitor their efficacy in ongoing trials. Conventional measures of therapeutic efficacy often rely upon measures of disability or cognition, or on imaging parameters such as atrophy, all of which may take several years to manifest (30). Changes in inflammation, or in metabolic processes within cells, will occur on a much more rapid time-scale and as such will be better biomarkers for clinical trials of novel therapies targeting these processes (31). Despite this, the technology to effectively monitor inflammation, especially within closed compartments such as the CNS, remains limited.

To provide the steps needed for the development of a clinical biomarker of early inflammatory activity, we require novel, safe, technologies capable of imaging longitudinal inflammation, a dynamic and multi-cellular process. Here, combined destructive (MALDI, histopathology, molecular biology) and translational (hyperpolarized MRI, plasma metabolomics) technologies to were used to readout changes in metabolism related to acute inflammatory challenge. Although our data are preliminary, these findings provide early proof-of-concept that hyperpolarized ^13^C approaches may contribute to such a biomarker pipeline (32). Understanding how the brain generates energy generally, not just in the context of neuroinflammation, is crucial given the number of diseases where metabolic disruption is an underlying pathological hallmark. Metabolism can be broadly divided into biosynthetic processes, which produce intermediates used to build cellular components, and energy-generating processes, which break down nutrients like glucose to produce ATP.

In addition to the metabolic alterations detected by hyperpolarized spectroscopy of the CNS, results demonstrated clear evidence of systemic metabolic reprogramming following acute neuroinflammatory challenge. Plasma metabolomics and lipidomics revealed a distinct separation between LPS-treated and control animals and identified significant shifts in metabolite and lipid abundance. The correlations between circulating lactate/pyruvate and specific plasma metabolites and lipid species further suggest that CNS inflammation induces a coordinated metabolic response across central and peripheral compartments. This systemic profile is consistent with established metabolic responses to innate immune activation, where inflammatory signalling drives increased glycolytic flux, altered TCA-cycle activity, and enhanced lipid mediator synthesis (31, 33). The plasma mass-spectrometry data therefore provide an external, easily accessible signature of the metabolic state induced by neuroinflammation and serve to validate the shifts in CNS pyruvate metabolism detected by hyperpolarized MRI. These observations should be interpreted cautiously given the limited sample size, but they reinforce the central aim of this feasibility study: to show that multimodal metabolic profiling can reveal coordinated CNS–peripheral responses to acute inflammation.

Importantly, the observation that peripheral metabolic alterations accompany early CNS metabolic changes supports the idea that hyperpolarized MRI detects biologically meaningful metabolic reprogramming, rather than non-specific changes in perfusion or tissue integrity. It also raises the possibility that combined peripheral metabolomics and metabolic imaging may offer a powerful multimodal biomarker framework for monitoring inflammatory activity *in vivo*. Future studies with larger cohorts, and using models with mixed pathology such as ischemia or multiple sclerosis, will be essential to validate these early observations.

Beyond the basic metabolic profiles, results demonstrated that there were changes in the one of the key enzymes that regulate entry of pyruvate into the TCA cycle, specifically in pyruvate dehydrogenase, with corresponding changes in mRNA expression levels of its regulators, pyruvate dehydrogenase kinases 1 and 2. These data, showing metabolic changes in the brain in response to LPS, are complementary to a previous study which utilised hyperpolarized MRI to assess chronic changes in brain metabolism where the authors showed an increase in [1-^13^C]lactate production in the brain at day 7 (10). This increase correlated with elevated Iba-1 and GFAP expression, and the authors postulate that the increase in lactate comes from the increased number of astrocytes and microglia. However, Le Page et al. did not aim to detect ^13^C bicarbonate as a co-polarized solution of [1-^13^C]pyruvate and ^13^C urea was infused, which may have obscured the ^13^C bicarbonate peak. Here we have demonstrated, at an earlier time point, that changes in ^13^C bicarbonate due to LPS challenge are detectable just 24 hours after insult, before meaningful changes in lactate formation are observed. These results thus provide early evidence that mitochondrial dysfunction may precede glycolytic shifts during acute neuroinflammation, offering new insight into the temporal sequence of metabolic changes in the brain.

Despite the fact that we are coming up to 90 years since the discovery of the citric acid cycle, it is only within the last 15 years that its metabolic intermediates have started to be observed *in vivo* in the brain (34). Our clinical capacity to rapidly image cellular processes is currently limited. Whilst MR spectroscopy can be used to measure specific cellular markers, such as glutamate or creatine, the ability to track dynamic changes in metabolism is often challenging and metabolites are required to be in the millimolar range to be resolved. Imaging inflammation using PET is feasible and many ligands show significant changes in pathology. However, the repeated exposure to radiation means this technology is not best suited to longtiduinal studies in chronic disease, such as those that might be required to study the development of post-stroke dementia, for example. By using hyperpolarised ^13^C labelled metabolites that readout on the TCA cycle, we have more of a handle on the downstream processes occurring within the CNS in response to injury and disease (5, 7, 10).

Whilst using hyperpolarized MRI as a tool for early detection of inflammatory activity may not currently be practical in the clinical setting, due to the often-late presentation of patients, it does point to the ability of the technology to provide a direct readout of druggable targets in the pre-clinical development and testing pipeline. To detect chronic immune activity in the brain, through the increased production of lactate, may be a more realistic use of the technology in neuroinflammatory disease settings where there are, otherwise, a lack of structural or functional changes the brain. By using static and dynamic means of measuring metabolism we have demonstrated that inflammation affects apparent CNS metabolism, as well as demonstrating the feasibility of using changes in metabolism as a proxy for measuring neuroinflammatory processes in the brain.

## Limitations and Conclusion

This study does have its limitations. Given the pilot-style feasibility nature of this work and the small sample size, some analyses were underpowered and results should be interpreted accordingly. The rodent model of neuroinflammation is somewhat reductionist and does not necessarily recapitulate the complexities of human disease. However, we have demonstrated that metabolic imaging in particular is translatable in a model of stroke (32) and our ongoing research using chronic models of stroke and multiple sclerosis aim to introduce additional levels of pathological complexity. Our main aim with this preliminary work was to demonstrate the feasibility of imaging the brain using hyperpolarized pyruvate after an inflammatory challenge and to provide evidence that this signal (supported by complementary spatial metabolomics and molecular biology *ex vivo*) can be used as a proxy for imaging ongoing inflammation *in vivo*. Our data builds on previous published work (10) and confirms hyperpolarized metabolic imaging is an effective way to monitor CNS inflammation.

## Author Declarations

### Ethics approval

All procedures conformed to the Animal (Scientific Procedures) Act (1986) and were approved by the UK Home Office and the University of Oxford Animal Ethics Committee (LERP and ACER, University of Oxford). All work was carried out under license number PP7444704 (Couch: Molecular Mechanisms of CNS Injury).

### Consent for publication

Not applicable

### Data availability

The datasets used and/or analysed during the current study are available from the corresponding author on reasonable request.

### Author contributions

JTG, AMD & YC conceptualised and designed the experiments; JTG, IE, DO and YC carried out the experiments; JTG, IE, DO, SN and YC analysed the experiments; all authors contributed to the writing and editing of the manuscript, all authors read and approved the final manuscript.

## Acknowledgements

The authors would like to acknowledge Nneoma Akinaro-Ejim, Vicky Ball and Aaron Axford, who provided technical support for the metabolic imaging. Additionally, the authors thank Matilda Kråkström for processing the mass spectrometric data. Mass spectrometry analysis was performed at the Turku Metabolomics Centre with the support of Biocenter Finland

